# Identification of a novel link connecting indole-3-acetamide with abscisic acid biosynthesis and signaling

**DOI:** 10.1101/2025.08.15.670611

**Authors:** José Moya-Cuevas, Paloma Ortiz-García, Adrián González Ortega-Villaizán, Irene Viguera-Leza, Andrés Pérez-González, Javier Paz-Ares, Carlos Alonso-Blanco, Jesús Vicente-Carbajosa, Stephan Pollmann

## Abstract

Plants regulate their developmental programs and their responses to environmental changes through a complex network of small signaling compounds, known as phytohormones. The role of auxins in promoting plant growth has been extensively investigated. Furthermore, previous studies have demonstrated that the accumulation of indole-3-acetamide (IAM), an auxin precursor, results in the suppression of plant growth, particularly primary root elongation. This observation led to the hypothesis that IAM or an IAM derivative exerts negative growth regulatory effects. However, the molecular mechanism by which IAM inhibits plant growth remains largely unelucidated. To gain deeper insight into the molecular mode of action of IAM, we conducted a comprehensive genome-wide association study (GWAS) using a highly diverse collection of 166 wild Arabidopsis accessions from the Iberian Peninsula. Consequently, we identified several genomic regions associated with a reduced response to IAM under controlled *in vitro* conditions, which included *ABA3* and *GA2ox2* as candidate genes. Sequence analyses, transcriptomics studies, and comparison of three-dimensional models generated for *ABA3* proteins encoded by the two major natural alleles identified in the collection of wild accessions suggested that IAM-triggered inhibition of primary root elongation is closely associated with the formation of abscisic acid (ABA) in *Arabidopsis thaliana* seedlings. Finally, physiological characterization of mutants for those candidate genes further corroborated that IAM activates ABA signaling. Our results demonstrate that IAM is intricately linked with ABA biosynthesis and signaling, thereby elucidating a novel node in plant hormone crosstalk.

## Introduction

For the implementation of intrinsic growth programs, as well as for the adaptation to their ever-changing environments, plants utilize a limited number of signaling molecules, commonly referred to as plant hormones. These are defined as naturally occurring substances that function at sub-micromolar concentrations to drive physiological processes either locally or in distant tissues (Santner *et al*., 2009; Davies, 2010). The various plant hormone groups regulate distinct subsets of biological processes independently. However, in recent years, accumulating evidence has led to the widely accepted understanding that the overall outcome of plant hormone action is determined by the extensive combinatorial activity of the different phytohormones and their cellular crosstalk through the spatiotemporal integration of their signaling pathways (Wolters & Jürgens, 2009; Depuydt & Hardtke, 2011; Ortiz-García *et al*., 2023; Wong *et al*., 2023). Plant hormones influence the development of a responding tissue by inducing transcriptional reprogramming of the corresponding cells. Transcriptional alteration can, consequently, result in physiological and metabolic adjustments (Hsu *et al*., 2021; Yoshida & Fernie, 2023). These initiated responses are additionally dependent on the specific properties of the responding tissues regarding their sensitivity and responsiveness to the given interacting signaling compound classes (Bradford & Trewavas, 1994; Knight & Knight, 2001).

Among plant growth-promoting phytohormones, indole-3-acetic acid (IAA), the most important endogenous auxin, plays a pivotal role in numerous plant growth and developmental processes, including cell division and expansion, meristem maintenance, differentiation, tropisms, apical dominance, senescence, leaf abscission, and flowering (Woodward & Bartel, 2005; Teale *et al*., 2006). Recent research has demonstrated the importance of stringent spatiotemporal regulation of auxin biosynthesis and its correlation with the aforementioned processes (Zhao, 2010; Xu *et al*., 2018). The primary precursor for IAA biosynthesis in plants is L-tryptophan (L-Trp). The indole-3-pyruvic acid (IPyA) pathway constitutes the principal route for IAA formation. This pathway proceeds from L-Trp through IPyA to IAA and involves the tryptophan aminotransferases TRYPTOPHAN AMINOTRANSFERASE OF ARABIDOSIS 1 (TAA1) and TRYPTOPHAN AMINOTRANSFERASE RELATED 2 (TAR2). Several alternative pathways, including the tryptamine pathway, the indole-3-acetaldoxime pathway, and the indole-3-acetamide (IAM) pathway, are also considered to contribute to IAA biosynthesis under specific conditions (Pollmann *et al*., 2006; Kasahara, 2016).

The IAM pathway comprises two enzymes: a tryptophan 2-monooxygenase (tms1, iaaM, aux1) that catalyzes the conversion of L-Trp to IAM, and an IAM-specific amidohydrolase (tms2, iaaH, aux2), which subsequently hydrolyzes IAM into IAA. Initially, this pathway was believed to occur exclusively in certain phytopathogenic bacteria of the genera *Agrobacterium*, *Pantoea*, and *Pseudomonas*. However, this understanding changed with the identification and characterization of the first IAM-specific amidohydrolases of Arabidopsis (*At*AMI1), tobacco (*Nt*AMI1), and several other plant species (Pollmann *et al*., 2003; Nemoto *et al*., 2009; Sánchez-Parra *et al*., 2014). More recent research has demonstrated that *ami1* T-DNA insertion mutant seedlings exhibit hypersensitivity to osmotic stress, and it has been proposed that the transcriptional suppression of *AMI1* expression by abiotic stress signals serves as a primary adaptation mechanism to this type of stress (Moya-Cuevas *et al*., 2021; Pérez-Alonso *et al*., 2021). Previous research has indicated a possible regulatory relationship between IAM and the biosynthesis of ABA. However, natural variation has not been utilized as a methodological approach to investigate the role of IAM as a potential signaling molecule, distinct from IAA.

In this study, we present the findings of a genome-wide association analysis aimed at identifying loci linked to the inhibitory effect of IAM on primary root growth, utilizing 166 Arabidopsis ecotypes from the Iberian Peninsula. Genetic analysis of a number of potential candidate genes indicated that *ABA3* is involved in the IAM-triggered negative regulation of primary root growth. Additionally, an investigation into the networks of IAM and ABA co-regulated genes uncovered a putative core response module that plays a role in managing responses to water and osmotic stress in Arabidopsis.

## Results

### IAM treatments highlight multiple genomic regions and candidate genes

To identify novel molecular components involved in IAM-dependent repression of primary root growth, a set of 235 Arabidopsis accessions from the Iberian Peninsula was utilized to conduct a genome-wide association study (GWAS). Following synchronization of all accessions, 166 natural accessions were selected for mock and IAM treatments and subsequent phenotypic analysis. As illustrated in **Figure 1**, a substantial range of variability was observed among the accessions tested regarding their response to IAM in the growth medium. To standardize the responses across various accessions, each 6-well plate included Col-0 (Columbia) seedlings as a reference for normalization throughout the experiments conducted. The subsequent Shapiro-Wilks test confirmed the normal distribution of the data (*W* = 0.952, *p* = 2.21 × 10^-5^). Although IAM repressed root growth in the majority of accessions, a promotive effect was observed in 11% of the accessions (**Figure 1b**). Normalized root response data obtained after mock and IAM treatment of the different accessions (**Supporting Information Table S1**) were utilized to perform the GWAS analysis using the easyGWAS application (http://easygwas.biochem.mpg.de).

**Figure 1.**
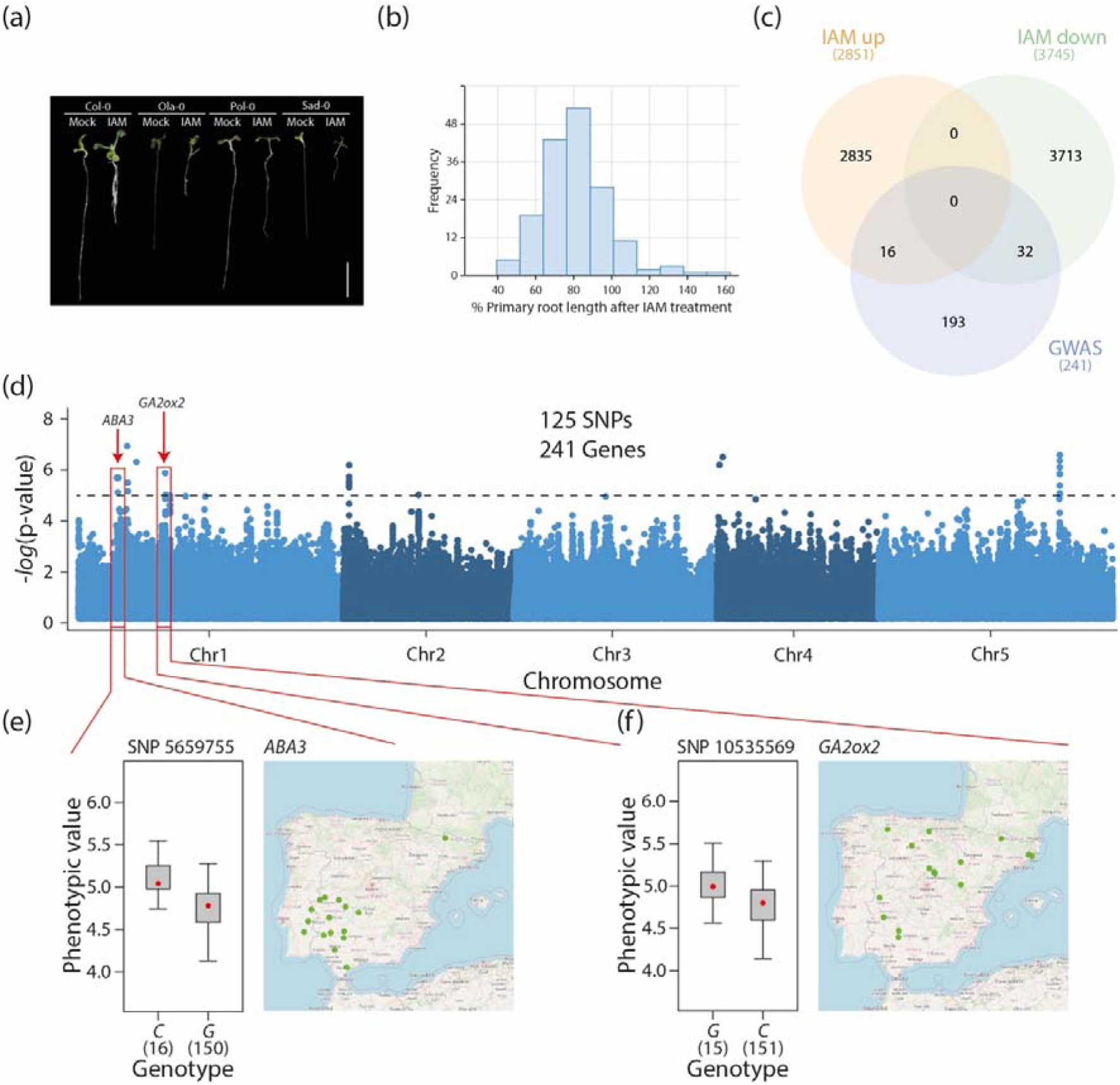
Arabidopsis accessions from the Iberian Peninsula display significant natural variations in their root growth response to IAM. (a) Representative images of four randomly selected accessions. (b) Frequency distribution of IAM-triggered primary root growth repression in 166 Arabidopsis accessions. (c) Venn diagram analysis of described IAM-responsive genes and the candidate genes identified through the GWAS approach. (d) Manhattan plots depicting IAM-induced primary root growth repression based on the linear mixed model. The dashed line represents -log10(p-values) > 5. (e,f) Normalized phenotypic values and geographic distribution of accessions exhibiting the minor frequency allele of SNP 5659755 (*ABA3*) and SNP 10535569 (*GA2ox2*). The box plots illustrate the median (red dot), the 25%-75% quartiles (grey box), and the extrema (whiskers) of the compared data.

To identify potential genomic regions associated with the tested phenotype, a relaxed *p*-value threshold (<10^-5^) was established (**Figure 1d**). Candidate genes located within a 10 kb region around the associated SNPs were compiled (**Supporting Information Table S2**). The SNP with the highest score (–log_10_(*p*-value) = 6.02) was SNP 16531216 on chromosome 5. Of the five genes located around the SNP, only two have been functionally characterized: *GLABRA 3* (At5g41315), which is related to trichome development (Bernhardt *et al*., 2003), and *UBIQUITIN CONJUGATING ENZYME 4* (At5g41340), which is involved in sugar metabolism and leaf senescence (Wang *et al*., 2022). For all SNPs that met the established criteria, a total of 241 possible target genes were obtained. Among the putative candidate genes, 22 transcription factor (TF) genes were identified, including three Lateral Organ Boundary (*LOB*) genes and two Agamous-Like (*AGL*) genes, as well as 12 development-related genes, including the secondary cell wall-related Arabidopsis NAM, ATAF1/2 and CUC2 (NAC) domain *NAC10* and the *HAIRY MERISTEM 3* (*HAM3*) gene. However, few of these genes exhibited documented expression in roots. The functional classification of the genes using the MapMan application (v3.6.0R1) provided additional evidence for a limited number of candidate genes related to plant hormones.

From the candidates presented in **Table 1**, the genes *ABA3* (At1g16540) and *GA2ox2* (At1g30040) were selected for further analysis due to their presence among the 16 and 31 GWAS candidate genes that have previously been reported to exhibit increased and decreased expression (**Figure 1c**), respectively, in response to a short-term IAM treatment (Ortiz-García *et al*., 2022).

**Table 1.** Functional classification of plant hormone-related genes detected by the GWAS analysis using MapMan.

| MapMan Bin code | AGI | Description | Significance threshold |
| --- | --- | --- | --- |
| 17.2.3 | At1g16510 | SAUR41 \| auxin-responsive family protein | 1.64E-05 |
| 17.3.3 | At3g61460 | BRH1 \| Brassinosteroid-responsive RING-H2 | 6.23E-05 |
| 17.6.1.11 | At5g51310 | gibberellin 20-oxidase-related | 5.23E-07 |
| 17.6.1.13 | At1g30040 | GA2ox2 \| GIBBERELLIN 2-OXIDASE | 4.5E-06 |
| 17.6.3 | At5g15230 | GASA4 \| GAST1 PROTEIN HOMOLOG 4 | 2.62E-05 |
| 17.1.1 | At1g16540 | ABA3 \| ABA DEFICIENT 3 | 1.64E-05 |

We identified 16 and 15 Arabidopsis accessions carrying the minor frequency allele for *ABA3* and *GA2ox2* candidate genes, respectively, which reduced the sensitivity to IAM (Figure **1e,f**). Analysis of the geographic distribution of these alleles revealed distinct patterns for both candidate genes. While *GA2ox2* alleles exhibited a dispersed distribution, the *ABA3* alternative allele was confined to a smaller geographic area in southwest Iberia (Figure **1e,f**). Notably, 13 out of the 16 accessions with the minor frequency allele of *ABA3* belong to the 14 accessions containing genetic group C4 previously described in the Iberian Peninsula (Tabas-Madrid *et al*., 2018), which has been previously characterized for a relevant adaptive trait dependent on ABA signaling, such as seed dormancy (Vidigal *et al*., 2016).

### Sequence analysis revealed strong linkage disequilibrium and structural polymorphisms in *ABA3*

To investigate the two selected candidate genes in more detail, we analyzed the quantity and distribution of genetic diversity in their genomic DNA sequences of 166 accessions. A total of 155 (95 in the promoter and 60 in the CDS) and 63 SNPs (37 in the promoter and 6 in the CDS) were segregating for *ABA3* and *GA2ox2*, respectively, corresponding to high and low nucleotide diversities (π-*ABA3* = 0.00415 (Promoter + CDS); π-*GA2ox2* = 0.00295 (Promoter + CDS)). As illustrated in **Figure 2**, SNPs were distributed throughout the length of the *ABA3* gene and exhibited strong linkage disequilibrium (LD), as the minor alleles were shared by the majority of the 16 Arabidopsis accessions with the minor allele detected by GWAS. Notably, 31 of the polymorphisms were in complete LD in these accessions, defining a highly differentiated allele referred to as ABA3-Alt (alternative), which encompassed 6 synonymous and 3 missense mutations: L102I, A150G, and L267V. In contrast, consistent with the low nucleotide diversity observed in *GA2ox2*, only two SNPs with partial LD distinguished the alternative allele identified as associated in GWAS analyses, and neither suggested an alteration in the primary amino acid sequence.

**Figure 2.**
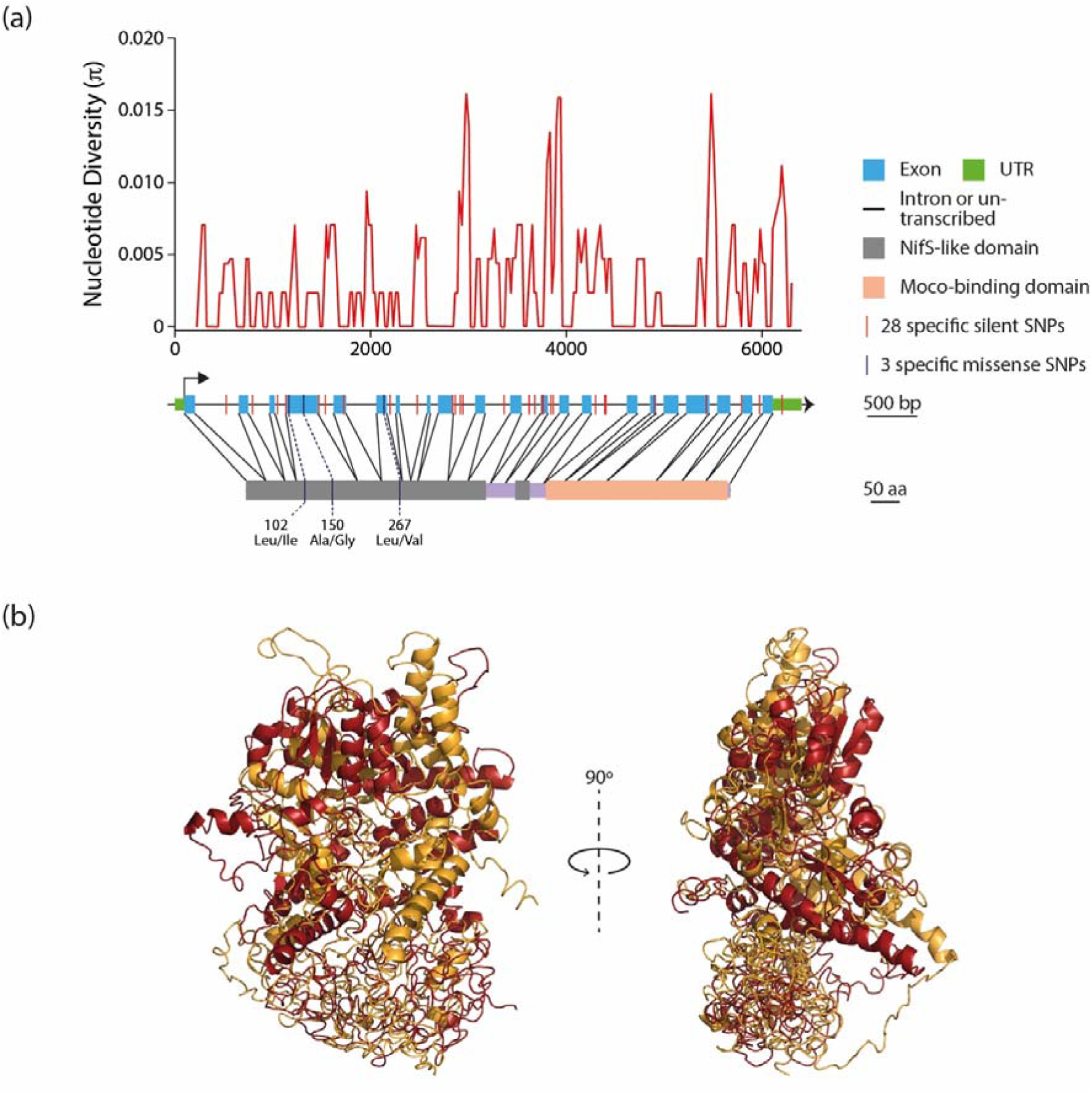
Analysis of the nucleotide diversity of *ABA3*. (a) Gene organization and nucleotide polymorphisms in *ABA3*. (b) Superimposition of the predicted three-dimensional structure models for ABA3_Ref (blue) and ABA3_Alt (red). The models have been inferred using the I-TASSER structural prediction server.

In order to assess the influence of missense mutations distinguishing the two alleles identified in *ABA3* on the functionality of the gene product, we conducted an analysis of the secondary and tertiary structures of the ABA3 protein as encoded by both the reference and alternative alleles. Although the three primary amino acid substitutions are conservative exchanges, the prediction of the secondary protein structures indicated minor differences in the length and organization of the α-helices, β-strands, and coiled regions (**Supporting Information Figure S1**).

Despite the high sequence identity of 99.64%, the reference ABA3 protein from Col-0 plants exhibits substantial differences from the alternative protein version found in the 16 Arabidopsis accessions. As illustrated in **Figure 2b**, the structural comparison of the homology-based protein models derived for ABA3_Ref and ABA3_Alt, obtained from the I-TASSER prediction server (Yang & Zhang, 2015) revealed that changes in the secondary structure of the two proteins encompass a considerable torsion of ABA3_Alt relative to ABA3_Ref. Such structural modifications can influence the geometry and steric properties of a protein. The activity and substrate affinity of enzymes are largely determined by optimal fixed distances between the amino acids of the catalytic center, as well as between the catalytic center and the cofactor binding sites. In this context, the L267V mutation, in particular, could interfere with the binding of the pyridoxal phosphate (PLP) cofactor to the conserved K271 residue in the NifS-like domain of the enzyme, which is essential for ABA3 activity (Heidenreich *et al*., 2005; Schwarz *et al*., 2009). Additionally, the mutations could affect protein stability. Therefore, the potential impact of the three mutations on protein stability was estimated using an integrative computational approach employing the DUET and the I-Mutant2.0 servers (Capriotti *et al*., 2005; Pires *et al*., 2014). While the ΔΔG values presented in **Table 2** showed minor discrepancies between the two tools, they consistently remained negative for all three mutations, indicating a predicted decrease in the stability of ABA3_Alt compared to ABA3_Ref.

**Table 2.**
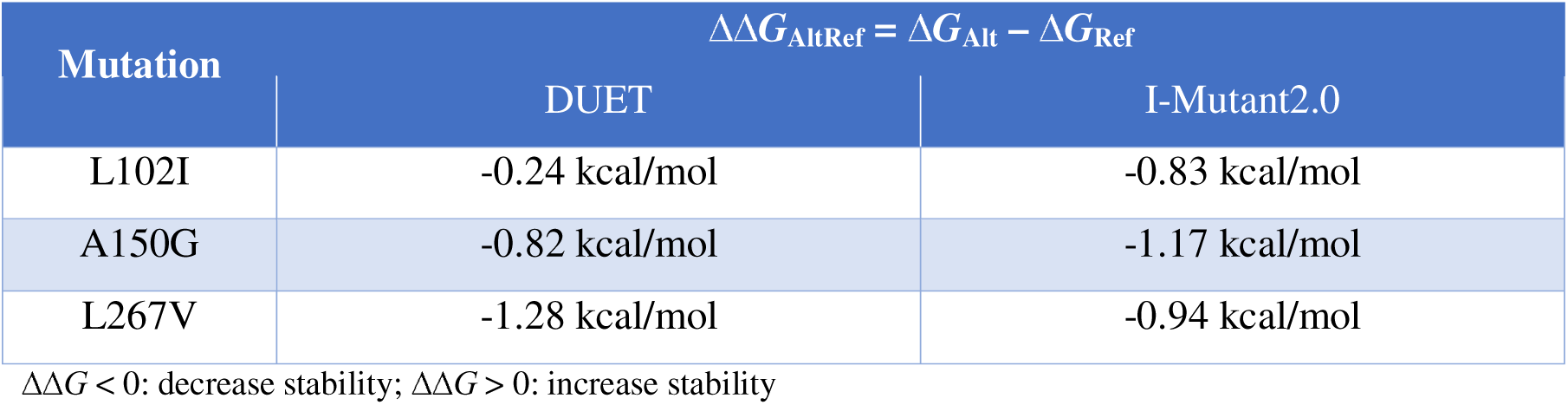
Calculation of Gibbs free energy of unfolding (Δ*G*) of ABA3 the modified protein variant found in the 16 identified Arabidopsis accessions compared to the reference protein from Col-0.

| Mutation | $\Delta\Delta G_{\text{AltRef}} = \Delta G_{\text{Alt}} - \Delta G_{\text{Ref}}$ | |
| --- | --- | --- |
|  | DUET | I-Mutant2.0 |
| L102I | -0.24 kcal/mol | -0.83 kcal/mol |
| A150G | -0.82 kcal/mol | -1.17 kcal/mol |
| L267V | -1.28 kcal/mol | -0.94 kcal/mol |

$\Delta\Delta G < 0$ : decrease stability; $\Delta\Delta G > 0$ : increase stability

### ABA3 affects the IAM triggered reduction of primary root growth

To further empirically investigate the role of *ABA3* and *GA2ox2* in the IAM-mediated growth repression, we obtained the corresponding knockout lines for both genes. The *GA2ox2* gene is member of a small isogene family of C19-GA2 oxidases comprising five members, *GA2ox1*, -*2*, *-3*, *-4*, and *-6*, which are reported to inactivate physiologically active GA through its oxidation (Rieu *et al*., 2008). In previous studies, we observed that not only *GA2ox2* but also *GA2ox6* is induced in the *ami1-2* mutant background that is impaired in the conversion of IAM into IAA (Pérez-Alonso *et al*., 2021). Furthermore, previous work identified a potential role of GA signaling in the response to IAM accumulation in the *ami1-2 rty1-1* double mutant (Sánchez-Parra *et al*., 2021). Consequently, we opted to incorporate the previously described *ga2ox1 ga2ox2 ga2ox3 ga2ox4 ga2ox6* quintuple mutant (*ga2ox q*) in the study to minimize the potential compensation of the *ga2ox2* mutation by the other isoforms.

ABA3 is a molybdenum cofactor sulfurase that is essential for the activation of aldehyde oxidases, such as ARABIDOPSIS ALDEHYDE OXIDASE 3 (AAO3), which catalyzes the final step in ABA biosynthesis (Seo *et al*., 2000; Bittner *et al*., 2001). For this reason, we decided to investigate the role of ABA biosynthesis and signaling in the processing of the IAM stimulus by analyzing the *aba3-1* and *abi5-7* knockout mutants, respectively.

First, we confirmed the previously described transcriptional regulation of *GA2ox2*, *GA2ox6*, and *ABA3* by IAM in the Col-0 reference ecotype (Ortiz-García *et al*., 2022). The gene expression levels in response to a short-term treatment with 10 µM IAM relative to mock-treated plants in wild-type Arabidopsis were analyzed by qPCR analysis. As demonstrated in **Figure 3a**, the three target genes exhibited moderate induction by IAM. The most pronounced upregulation was observed for *GA2ox2*, followed by *ABA3*. The effect of IAM on the expression of *GA2ox6* was minimal.

**Figure 3.**
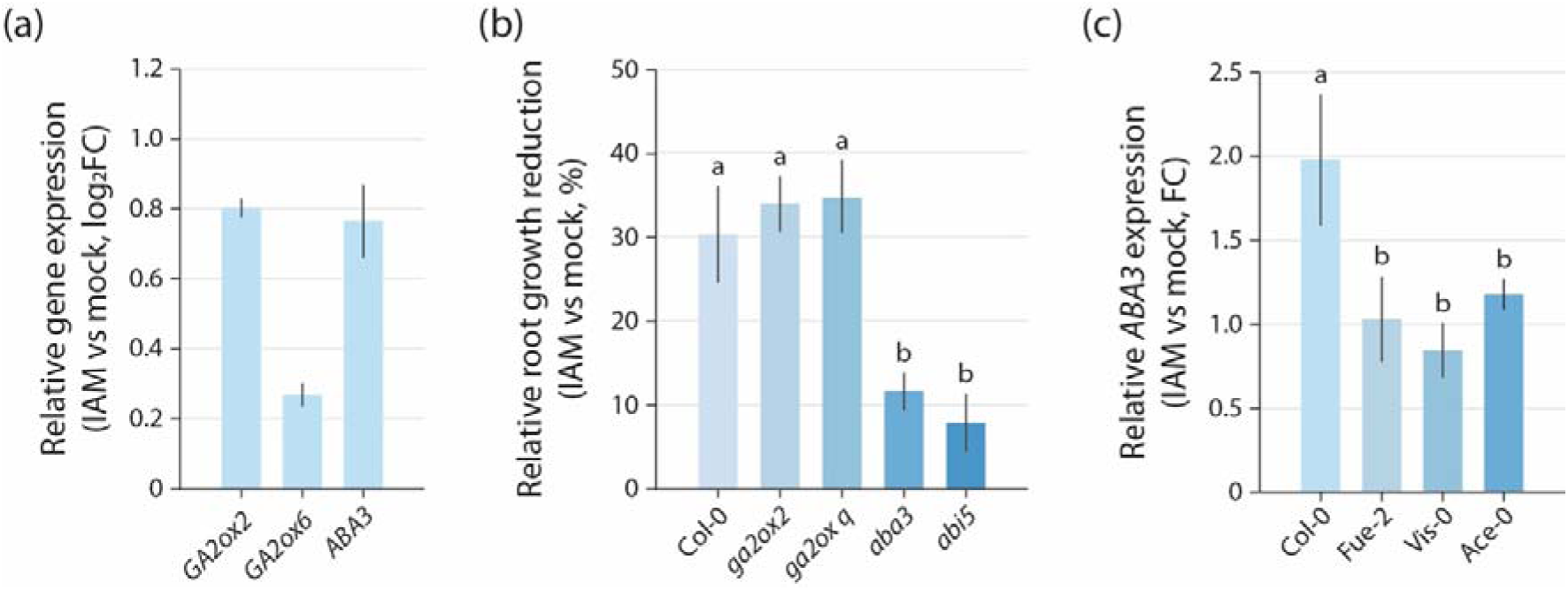
Transcriptomics and reverse genetics analysis of GA degradation and ABA biosynthesis and signaling related genes. (**a**) Quantification of *GA2ox2*, *GA2ox6*, and *ABA3* expression in Arabidopsis wild-type (Col-0) seedlings. The plot presents means ± SE (n=9) of the relative expression levels comparing the samples from mock treated plants with plants that were treated for 2 h with 10 µM IAM. (**b**) Comparative analysis of IAM-dependent primary root growth inhibition in GA degradation (*ga2ox2*, *ga2ox q*), an ABA biosynthesis (*aba3*), and ABA signaling (*abi5*) mutants. The values represent the means ± SE (n=24) of the root growth reduction relative to the length of wild-type (Col-0) Arabidopsis plants grown under control conditions. (**c**) IAM-mediated *ABA3* induction in three IAM insensitive Arabidopsis accessions compared to the Col-0 reference accession. For the statistical analysis, one-way ANOVA with a post hoc Tukey-Kramer test was employed. Different letters indicate significant differences between means (p ≤ 0.05).

The observed transcriptional responses indicated an enhancement in the catabolism of bioactive GA concomitant with the upregulation of ABA biosynthesis. Subsequently, we investigated the root growth response in various IAM- and mock-treated Arabidopsis mutants. As all mutant lines shared the same Col-0 background, this wild-type accession was utilized as a reference to evaluate the impact of the different mutations on IAM-dependent root growth inhibition. As illustrated in **Figure 3b**, Col-0 control seedlings exhibited approximately 30% shorter primary roots when treated with IAM. For the *ga2ox2* and *ga2ox q* mutants, a 34% and 35% reduction of primary root growth in IAM-treated seedlings was observed, respectively. However, these latter values of root growth reduction were not significantly different from the wild-type control. In contrast, ABA biosynthesis and signaling mutants demonstrated a significantly lower inhibition of root growth triggered by IAM. For the *aba3* mutant, only a 12% reduction of primary root growth was recorded, while the ABA signaling mutant, *abi5*, exhibited only an 8% reduction. It is noteworthy that the *aba3* and *abi5* mutations had a similar significant effect on IAM-triggered primary root growth inhibition, thus suggesting that ABA synthesis and signaling are equally important for this phenotype. Collectively, these findings indicate that IAM-induced ABA biosynthesis and signaling events contribute to the inhibition of IAM-dependent root growth inhibition observed in wild-type Col-0 control plants, while C19-GA2 oxidases are likely not involved in the generation of the root growth response to IAM. Subsequently, we investigated whether the transcriptional regulation of *ABA3* gene expression differed between the Col-0 plants and plants from three selected representative IAM-insensitive Arabidopsis accessions (**Figure 3c**). qPCR analysis revealed a significant difference in *ABA3* induction in the accessions Fue-2, Vis-0, and Ace-0 compared to Col-0. In contrast to Col-0, which exhibited the anticipated induction of *ABA3* following short-term treatment with IAM, the three selected accessions did not demonstrate such induction. This observation indicates that the insensitivity to IAM is largely attributable to modifications in the transcriptional regulation of *ABA3*.

### IAM triggers ABA signaling in Arabidopsis roots

Given the substantial impairment of ABA biosynthesis and signaling mutants in the response to IAM and the previously documented impact of endogenous IAM accumulation on cellular ABA levels (Pérez-Alonso *et al*., 2021), we postulated that IAM application induces ABA synthesis and subsequent signaling in Arabidopsis roots. To examine this hypothesis, alterations in ABA signaling were assessed in ABA signaling reporter lines following a short-term treatment with 20 µM IAM. As illustrated in **Figure 4a**, the exposure of the reporter lines to IAM elicited an ABA signaling response comparable to that observed in the ABA-treated control samples.

**Figure 4.**
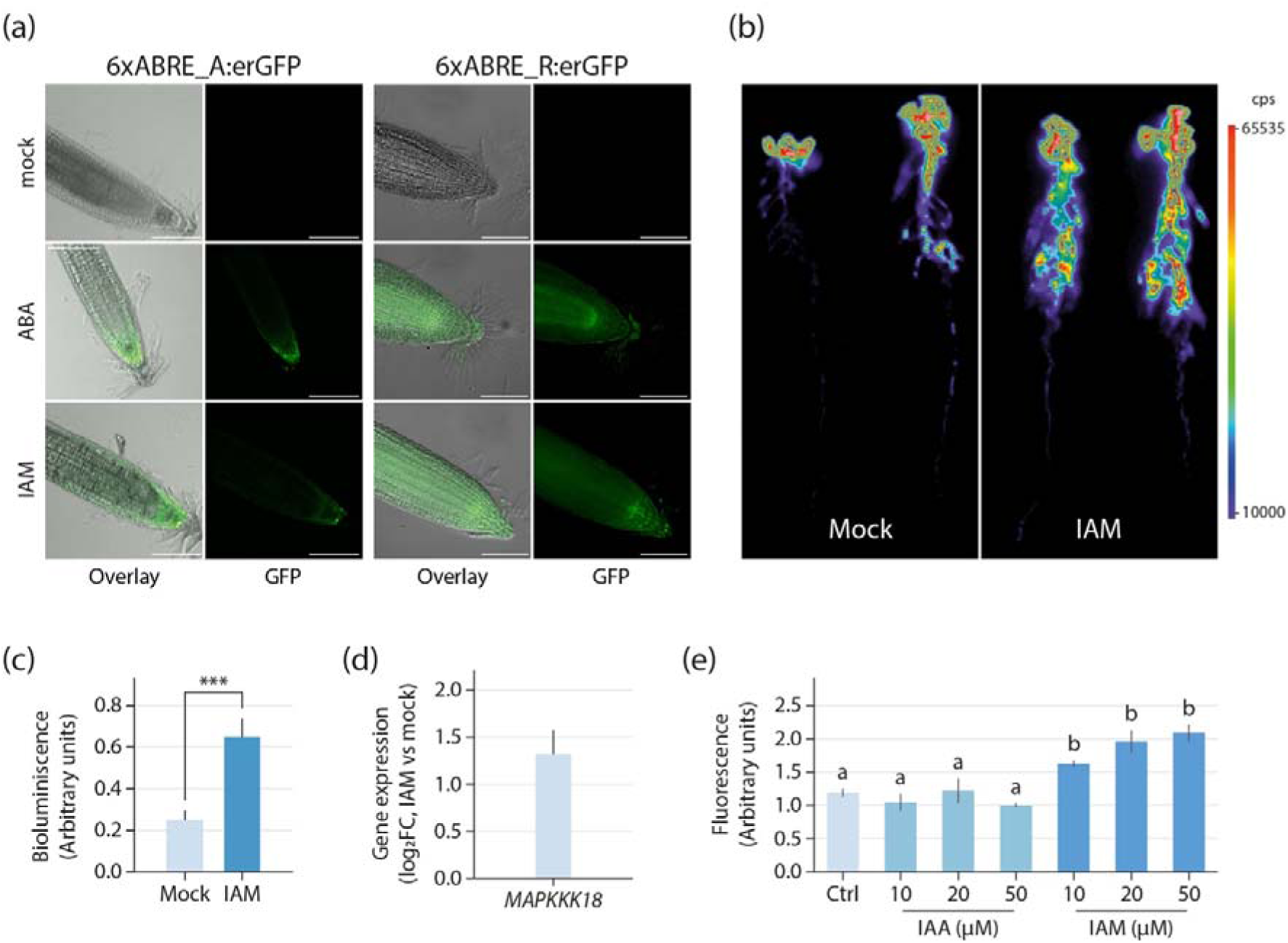
IAM induced ABA signaling. (**a**) Representative images of the effect of a 2 h treatment with 10 µM ABA and 20 µM IAM in comparison to mock controls monitored by using two synthetic promoter reporter lines for ABA signaling. Scale bar = 100 µm. (**b**) Confirmation of IAM triggered ABA signaling at the whole plant level using the pMAPKKK18::Luc^+^ bioluminescence promoter reporter line. The color scale gives the bioluminescence value. (**c**) Quantification of the average bioluminescence signal in mock and IAM treated reporter plants. The graph depicts means ± SE (n = 5). Student’s *t*-test: \*\*\**p* ≤ 0.001. (**d**) Quantification of transcriptional alterations of the *MAPKKK18* (At1g05100) gene in response to a 2 h treatment with 20 µM IAM by qRT-PCR. The bar shows the mean ± SE of n = 3 independent measurements. (**e**) Quantification of the average fluorescence signal in synthetic ABA reporter lines following mock treatment or the application of increasing concentrations of IAA and IAM, as indicated in the image. The graph illustrates the means ± SE (n = 5). Statistical significance was determined using one-way ANOVA with a post hoc Tukey-Kramer test. Different letters indicate significant differences between means (*p* ≤ 0.05).

Furthermore, we investigated the effect of IAM treatment at the whole plant level. To this end, we examined the effect of IAM treatment on ABA signaling using the pMAPKKK18::Luc^+^ reporter line, which confirmed the promotion of ABA signaling (**Figure 4b**). The bioluminescence signal was quantified under these conditions (**Figure 4c**), and the induction of the *MAPKKK18* gene by IAM was confirmed through qRT-PCR analysis (**Figure 4d**). It is noteworthy that the observed effect was predominantly localized to the roots.

Subsequently, we investigated whether ABA signaling is specifically triggered by IAM or if IAA could also induce ABA signaling in the 6xABRE_A::erGFP and 6xABRE_R::erGFP reporter lines. To address this question, the reporter lines were either mock-treated or subjected to a 2 h short-term treatment with IAA and IAM, respectively, at increasing concentrations (**Figure 4e**). The quantification of the fluorescence provided evidence for a statistically significant increase in ABA signaling induction when the reporter lines were treated with IAM. Conversely, the IAA treatment had no significant effect on ABA signaling, suggesting an IAM-specific effect.

### IAM and ABA regulate a common transcriptional response module

Considering the likely crosstalk between IAM and ABA, we subsequently examined whether convergent gene regulatory networks could be inferred by comparing existing transcriptomics data sets. To this end, a data set on IAM-triggered transcriptional responses in the *ami1* null mutant *ami1-2* (Pérez-Alonso *et al*., 2021) was compared with publicly available data (GSE39384) deposited in the gene expression omnibus (GEO) database (Barrett *et al*., 2012) regarding the response of Arabidopsis seedlings to ABA treatment (Goda *et al*., 2008) (**Supporting Information Table S3**). As illustrated in **Figure 5**, a shared group of 206 differentially expressed genes (DEGs) (8.3% of the total compared genes) was identified in the intersect between the 1,020 and 1,673 DEGs in response to 3 h IAM or ABA treatments, respectively (cutoff of log_2_(fold change) ≥ |1|, FDR ≤ 0.05).

**Figure 5.**
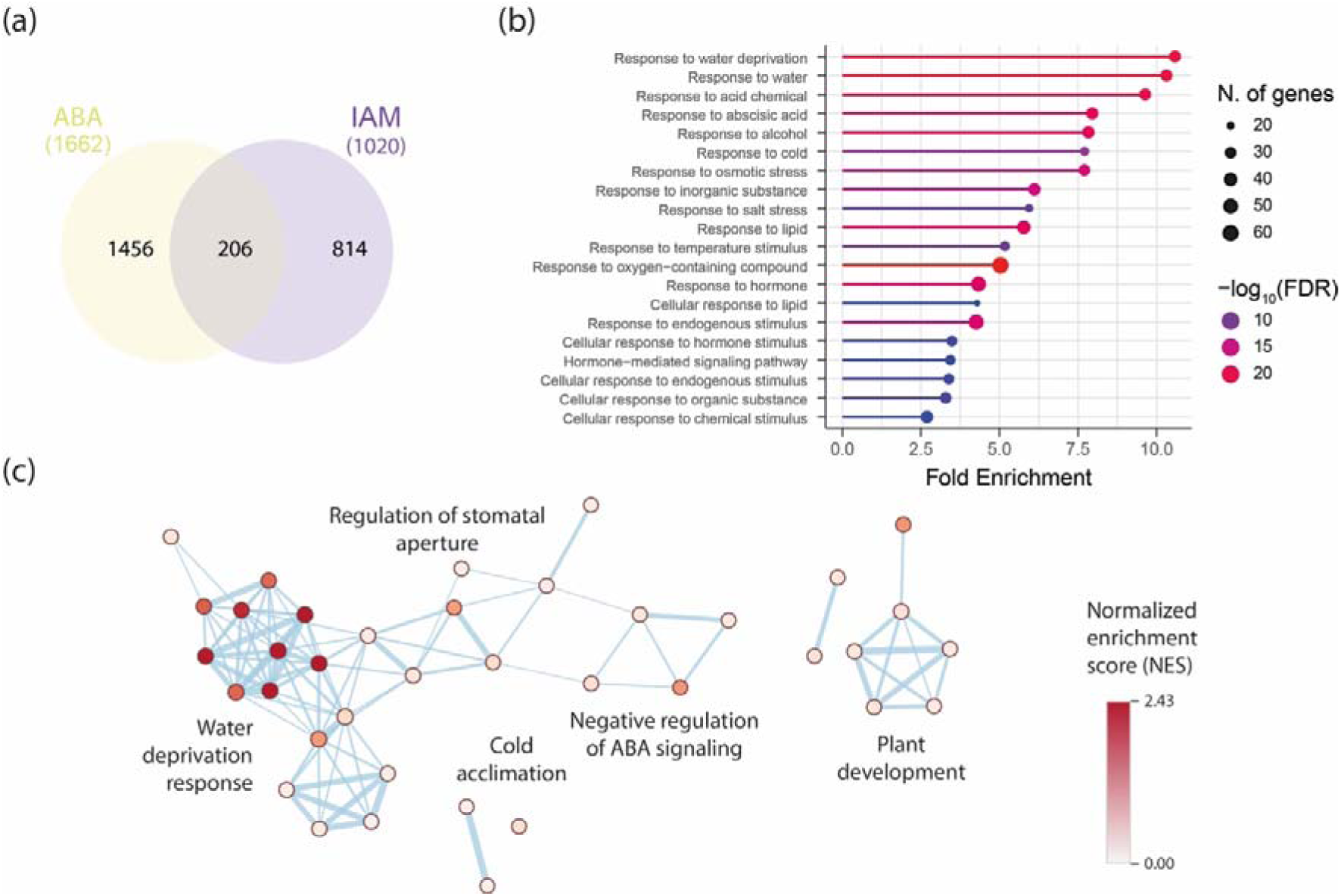
Comparison of IAM and ABA regulated genes in Arabidopsis. (**a**) Venn diagram analysis showing the numbers of DEGs in response to a 3 h IAM and ABA treatment. (**b**) GO enrichment analysis of the 206 genes in the intersection between IAM and ABA regulated DEGs. Each circle in the figure represents a distinct GO term, and the circle size indicates the number of genes enriched in the corresponding GO term. The significance of the observed gene enrichment is represented by a color gradient referring to the -log_10_(FDR). (**c**) GO term enrichment map for the 31 DEGs in the core *RD29B*/*HAI1* network. GO terms that share members are shown in connected clusters. Cluster labels were retrieved using the AutoAnnotate v1.4.1 application in Cytoscape. The node color intensity reflects the normalized enrichment score (NES).

To further characterize the physiological function(s) of genes shared by both ABA and IAM responses, we conducted a gene ontology (GO) analysis and a functional network analysis. A comprehensive analysis of the GO classification of the selected DEGs revealed a potential enrichment of genes in biological processes associated with numerous stress responses (**Figure 5b, Supporting Information Table S3**). In addition to an anticipated enrichment of general hormone response-related GO terms, such as GO:0009725 (response to hormone), GO:0032870 (response to hormone stimulus), and GO:0009755 (hormone-mediated signaling pathway), and a directly ABA-related GO term, i.e. GO:0009737 (response to abscisic acid), we identified several abiotic stress-related classifications among the significantly enriched GO terms. The GO classification with the highest enrichment was GO:0009414 (response to water deprivation), followed by GO:0006970 (response to osmotic stress), GO:0009409 (response to cold), and GO:0009651 (response to salt). These GO terms indicate the involvement of the IAM and ABA regulated target genes in the adjustment of the cellular osmotic or solute potential (ΨS).

Subsequently, a functional association network analysis was conducted using the stringApp in Cytoscape for the 206 selected genes, resulting in a network with 206 nodes and 245 edges. Analysis of the network topology identified the genes *RESPONSIVE TO DESICCATION 29B* (*RD29B*) and *HIGHLY ABA INDUCED PP2C GENE 1* (*HAI1*) as central components in the network. *RD29B* exhibited the highest degree of connectivity with 22 connections, while *HAI1* demonstrated the highest betweenness centrality of the core network. **Figure 5c** illustrates the subnetwork inferred from *RD29B* and *HAI1* and their 29 directly connected genes, including the PP2C genes *ABA-INSENSITIVE 1* (*ABI1*), *ABI2*, *HYPERSENSITIVE TO ABA 1* (*HAB1*), and *HAI2*. Moreover, *RD29B* and *HAI1* appeared to be functionally associated with a series of TFs, including *HB-7* and *HB-12*, two Homeodomain-Leucine Zipper subfamily I (HD-ZIP I) TFs (Ré *et al*., 2014), the NAM, ATAF1/2 and CUC2 (NAC) family TFs *NAC019* (Sukiran *et al*., 2019), *NAC032* (Maki *et al*., 2019), and *NAC092|NAC6|ORE1* (Escamez *et al*., 2020), as well as the MYELOBLASTOSIS (MYB) family TFs *MYB74* (Ortiz-García *et al*., 2022). Subsequently, the 31 genes in the *RD29B/HAI1* subnetwork were subjected to an enrichment analysis using the Enrichment Map application. The nodes in **Figure 5c** represent biological processes, and the edges represent gene crosstalk between the different processes (calculated using *p* < 0.05, FDR < 0.1, and the Jaccard coefficient cutoff of 0.25), which links biological processes that share a substantial proportion of genes, thus reducing the redundancy that exists in GO databases. In summary, biological processes related to the response to water deprivation, including adjustment of the cellular osmotic potential and plant growth, are the most prominent physiological functions regulated by the overlapping action of IAM and ABA.

## Discussion

The regulation of root development is highly intricate and essential for plant fitness (Motte *et al*., 2019). The root system anchors the plant in the soil and serves as the organ responsible for the absorption of water and inorganic nutrients (Robe & Barberon, 2023). Regarding the latter, roots function as source tissues from which nutrients are subsequently distributed to sink tissues, including the photosynthetically active aerial plant organs, in a meticulously regulated manner (Chang & Zhu, 2017).

Our previous studies highlighted a growth-inhibiting effect of IAM on primary roots. Mutations in the *AMI1* gene caused an accumulation of IAM in the seedlings and led to a substantial change in the root phenotype characterized by shorter roots and a reduced root branching density. In addition, IAM treatments of *ami1* knockout and conditional *AMI1* overexpression mutants resulted in longer and shorter primary roots, respectively, which was partly attributed to the conversion of IAM to IAA by IAM-specific amidases (Pérez-Alonso *et al*., 2021). By contrast, another study put further emphasis on the growth inhibiting effect of IAM on Arabidopsis seedlings. With the objective to rescue mutants in the *SUPERROOT 1*|*ROOTY* gene, the *rty1-1* mutant was crossed with the *ami1-2* mutant to block the increased conversion of IAM to IAA in this mutant (Sugawara *et al*., 2009). However, the *ami1-2 rty1-1* double mutant showed an unexpected growth arrest resulting in non-viable seedlings, which was traced back to a massive accumulation of IAM in the double mutant (Sánchez-Parra *et al*., 2021).

In this study, we employed a GWAS approach to advance our understanding of the role of IAM in regulating the development of the root system. By utilizing natural variations as a molecular tool, we aimed to identify novel components associated with IAM-mediated root growth inhibition. Phenotypic analysis of 166 Arabidopsis accessions from an Iberian Peninsula collection (Arteaga *et al*., 2021) revealed substantial variation in the root response to IAM in the growth medium, which varied between hypersensitive accessions, such as Ria-0 with a nearly 65% reduction of primary root length, and hyposensitive accessions, such as Ace-0 that showed no response to IAM or even growth induction relative to the control (**Figure 1**). Notably, among the 241 genes with significant associations, only six were found to be directly related to plant hormones (**Table 1**). We further investigated two of these genes, *ABA3* and *GA2ox2*, as they have been previously shown to play a role in the differential responses to IAM across various genotypes (Ortiz-García *et al*., 2022). In the case of *GA2ox2*, the most significantly associated SNP 10535569 was found 1888 bps upstream of the gene, which shows no strong LD with SNPs within the coding sequence of the gene. Moreover, the analysis of the individual and quintuple *ga2ox* mutants failed to show any implication of GA2ox proteins in IAM-mediated root growth inhibition. On the contrary, for the *ABA3* gene, the minor frequency allele at the most significant SNP in position 5659755 was shared by 16 accessions. This SNP was in complete LD with numerous SNPs in the coding sequence, including three missense mutations, which led us to compare the predicted three-dimensional structure of ABA3 protein between the shared alternative variant (ABA3_Alt) and the Col-0 reference version (ABA3_Ref). This analysis suggested substantial structural changes in the ABA3 protein encoded by the specific Iberian allele (Figure **2**). Most intriguing, however, was the observation that some representative IAM-insensitive accessions showed no transcriptional response to IAM, which is likely to be attributed to the considerably high nucleotide diversity of π-*ABA3* = 0.0073 detected for the promoter region of the 16 identified accessions (**Figure 3c**). Furthermore, accessions with this allele belong to an Iberian specific genetic lineage that is distributed exclusively in South-western Iberia (**Figure 1**), a geographic region characterized by high average temperature and low precipitation (Brenan et al., 2014). This climatic distribution has been previously shown to likely impact the life-cycle phenology of Arabidopsis populations belonging to this genetic lineage, as supported by the positive correlation between mean annual temperature and seed dormancy in this region (Vidigal *et al*., 2016; Marcer *et al*., 2018). Therefore, the *ABA3* allele of these accessions might contribute to adaptation to such environmental conditions, although, given the broad genetic differentiation of this lineage, we cannot discard that such adaptation is caused by other correlated genetic components (Picó *et al*., 2008; Brennan *et al*., 2014).

Given the observed insensitivity to IAM in the identified accessions possessing the minor frequency allele of ABA3, it can be hypothesized that the resultant ABA3_alt protein exhibits diminished enzymatic activity, leading to a significant reduction in ABA content. Although the proposed explanation is supported by the predicted reduced stability of the alternative protein, it does not align with the previously documented high seed dormancy, as indicated by DSDS50 (days of seed dry storage required to achieve 50% germination) values > 400 for the identified accessions (Vidigal *et al*., 2016). Seed dormancy is a crucial adaptive trait that optimizes seed germination in response to favorable environmental conditions. The regulation of seed dormancy and germination is predominantly governed by the intricate interaction between GA and ABA biosynthesis and signaling pathways (Liu & Hou, 2018). It is widely recognized that ABA serves as a significant positive regulator of seed dormancy (Finch-Savage & Leubner-Metzger, 2006). A deficiency in ABA during seed development leads to a lack of primary dormancy, whereas excessive ABA production during seed maturation enhances dormancy and typically results in delayed germination (Nambara & Marion-Poll, 2003). Consequently, a general reduction in ABA synthesis in accessions possessing the minor frequency allele of *ABA3* can be ruled out. This highlights the reduced or absent transcriptional response to IAM in the tested IAM-insensitive accessions. Consequently, it can be concluded that alterations in the transcriptional regulation of the *ABA3* gene by IAM are responsible for the abnormal root growth phenotype.

Given the significant reduction in IAM-dependent inhibition of primary root growth observed in the *aba3* ABA biosynthesis mutant and the *abi5* ABA signaling mutant (**Figure 3b**), we deduce that IAM-induced ABA synthesis, facilitated by the induction of *ABA3* (**Figure 3a**), is crucial for the repression of root growth in the reference Arabidopsis accession Col-0. The functional association between ABA3 and IAM-mediated root growth repression provides a link that integrates the IAM pathway of auxin biosynthesis with the synthesis of the stress hormone ABA. To empirically establish a connection between IAM and subsequent ABA signaling events, we hypothesized that IAM treatment should induce discernible changes in ABA signaling. In support of this hypothesis, the treatment of ABA signaling reporter lines with IAM demonstrated a substantial activation of ABA signaling and indicated that the effect of IAM was primarily confined to the roots (**Figure 4a,b**). Crucially, it was observed that increasing levels of IAM significantly induced ABA signaling, whereas similar experiments with IAA did not lead to a notable induction of ABA signaling. These results imply that IAM may have a function beyond its role as an intermediate in a secondary pathway of auxin biosynthesis, potentially acting as an independent signaling molecule (**Figure 4e**).

The physiological and molecular functions of ABA as a signaling molecule in plant responses to abiotic stressors are well established. An increase in cellular ABA levels has been convincingly linked to salt- (Jia *et al*., 2002; Holsteens *et al*., 2022), cold- (Shinkawa *et al*., 2013), and drought stress (Kuromori *et al*., 2018). At low concentrations (0.1 µM), ABA promotes root growth in a dose-dependent manner, whereas treatments with 1 µM ABA inhibit root growth (Ghassemian *et al*., 2000; Yoshida *et al*., 2019). The inhibitory effect of high ABA concentrations on root growth involves the suppression of cell division in the apical meristems and the repression of cell expansion in the root elongation zone (Takatsuka & Umeda, 2014; Yang *et al*., 2014). Moreover, the growth inhibitory effects are further mediated by components of the ABA signaling cascade. Several of the 14 ABA receptor molecules in Arabidopsis, such as PYR1, PYL1, PYL2, PYL4, PYL5, and PYL8, act redundantly, thus contributing to the inhibition of primary root growth (Park *et al*., 2009; Gonzalez-Guzman *et al*., 2012; Antoni *et al*., 2013). After ABA binding to the receptors, the negative co-regulators ABI1, ABI2, HAB1, and PP2CAs are sequestered by the receptors and thereby inactivated (Rubio *et al*., 2009; Thole *et al*., 2014). This liberates the sucrose non-fermenting 1 related protein kinases (SnRKs) SnRK2.2, SnRK2.3, and SnRK2.6 that act downstream of the PP2Cs and promote the ABA inhibition of primary root growth (Fujii *et al*., 2007; Zheng *et al*., 2010). The comparative transcriptomics analysis of DEGs between the IAM accumulating *ami1-2* mutant and ABA treatment underpinned a substantial overlap of the responses. More than 20% of the DEGs in the *ami1-2* mutant were also found to respond to ABA. The classification of the 206 DEGs in the intersection pointed towards an involvement in responses to drought, cold, and osmotic stress (**Figure 5**). A functional network analysis revealed a core subnetwork comprising a considerable number of PP2C type protein phosphatases, including *ABI1*, *ABI2*, *HAB1*, as well as *HAI1* and *HAI2*. The induction of these negative regulatory components possibly points to an involvement of IAM in the transcriptional control of a feedback loop that negatively regulates drought stress responses, such as the control of osmoregulatory solute accumulation, which involves the clade A PP2Cs HAI1, HAI2, and HAI3 (Bhaskara *et al*., 2012).

In summary, our research presents novel evidence supporting the involvement of *ABA3* in IAM-mediated primary root growth inhibition. Consistent with the previously documented induction of *NCED3* and the resultant elevated ABA levels in the *ami1* mutant alleles (Pérez-Alonso *et al*., 2021), the observed induction of *ABA3* further substantiates the proposed model illustrated in **Figure 6**. According to data extractable from the ePlant webtool (Waese *et al*., 2017), abiotic stress signals, such as cold, salt, and osmotic stress, exert a negative regulatory effect on the expression of *AMI1* (**Supporting Information Figure S2**). This regulation leads to increased levels of IAM, which in turn enhance ABA biosynthesis through the induction of *ABA3*, thereby attributing a signaling molecule role to IAM. Consequently, when *AMI1* expression is diminished in response to abiotic stimuli, an elevation in *ABA3* expression can be anticipated; a hypothesis that is predominantly validated by transcriptomic data accessible in public repositories (**Supporting Information Figure S2**).

**Figure 6.**
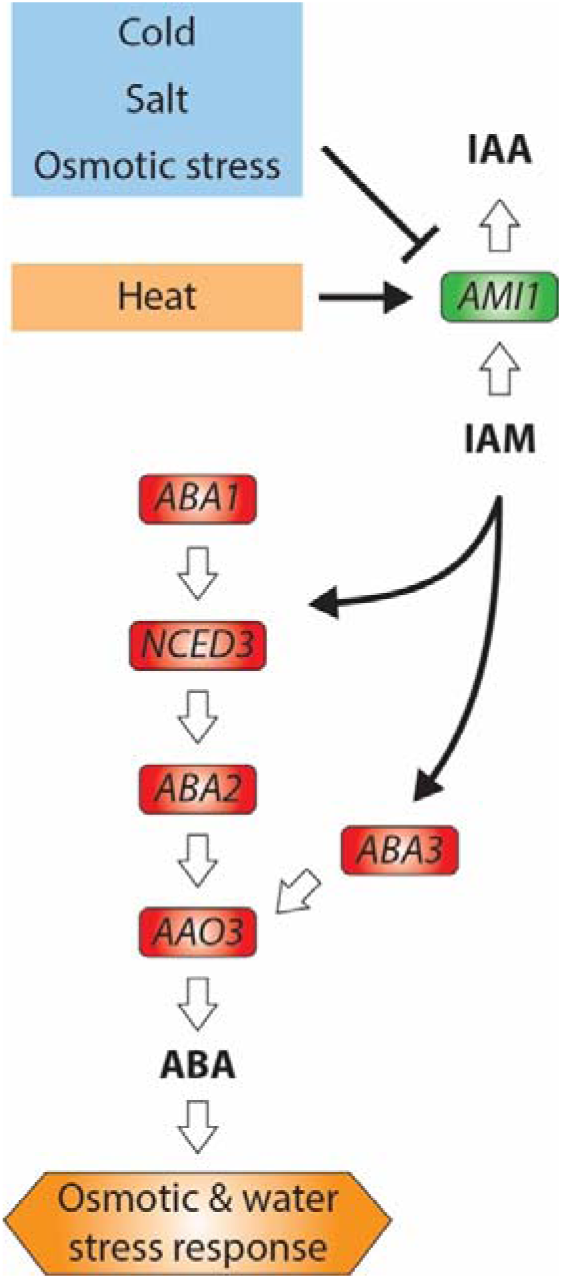
Proposed model for the regulation of abscisic acid (ABA) biosynthesis by indole-3-acetamide (IAM). Straight white arrows denote enzymatic processes, while black lines signify regulatory processes. Bold letters emphasize signaling molecules. Abiotic stress conditions regulate the expression of *AMI1*, consequently influencing the levels of IAM. The accumulation of IAM enhances the expression of ABA biosynthetic genes, including *NCED3* and *ABA3*. The upregulation of these genes results in elevated ABA levels and signaling, which subsequently governs downstream processes associated with osmotic and water stress responses.

Furthermore, our findings reveal a previously unrecognized shared stress response module that is likely implicated in the regulation of abiotic stress responses in Arabidopsis. Investigating the fundamental mechanism of this identified stress response module, which is proposed to integrate varying levels of IAM, represents an intriguing prospect for future research. In this context, a primary objective will be to investigate the existence of an IAM receptor and to identify the IAM-responsive transcription factor that presumably governs the transcriptional response of *ABA3* in Col-0, but not in the accessions carrying the minor frequency allele of *ABA3*.

## Experimental procedures

### Plant material

A previously published collection of 235 *Arabidopsis thaliana* accessions from the Iberian Peninsula (Arteaga *et al*., 2021) was synchronously grown to multiply the seed material. After the plants completed their life cycle, the seeds were harvested and stored under homogeneous conditions (22 [C, 50% RH). For the genome-wide association study (GWAS) experiment, accessions that showed low or null germination rates were discard. A resulting subset of 166 accessions was selected for the final experiment. The metadata of all the employed accessions are provided in **Supporting Information Table S1**. Each accession corresponds to the self-progeny of a randomly sampled individual plant per population. In addition, the following Arabidopsis accessions and mutants, respectively, were used: Col-0 (stock N1092), 6×ABRE_A::erGFP and 6×ABRE_R::erGFP (Wu *et al*., 2018), pMAPKKK18::Luc^+^ (García-Maquilón *et al*., 2021), *aba3-1* (Léon-Kloosterziel *et al*., 1996), *abi5-7* (Nambara *et al*., 2002), *ga2ox2-1* (Rieu *et al*., 2008), and *ga2ox q* nls GPS1 (Rizza *et al*., 2021).

### Root growth inhibition assay

Seeds were surface-sterilized, using sodium hypochlorite and 70% (v/v) ethanol, and then stratified at 4°C for two days. 15-20 seeds per accession were initially grown in 6-well plates, containing 10 ml per well of liquid 0.5× Murashige and Skoog (MS) media (Murashige & Skoog, 1962) supplemented with 1% (w/v) sucrose, and germinated for four days under controlled long-day growth conditions (22°C, 16 h light/8 h dark). Thereafter, the liquid media was replaced by 0.5× MS media containing only 0.1% (w/v) sucrose. Half of the seedlings were treated with 10 μM IAM (from 1 mM stock in ethanol), while the other half (control replicates) was mock treated with the corresponding volume of IAM-free ethanol. After 12 days, the seedlings were spread out on a solid support and photographs were taken to facilitate the measurement of the primary root length using the Fiji software (Schindelin *et al*., 2012). The primary root length of all tested accessions was normalized against the results obtained for Col-0 reference plants that were added to each 6-well plate.

### Genome-wide association study

The genome sequences and SNP data for the 166 Arabidopsis accessions used for GWAS are included in the 1001 Genomes project (Genomes Consortium, 2016) and in the publicly databases of the online applications for GWA Mapping in Arabidopsis, GWAPP (Seren *et al*., 2012) and easyGWAS (Grimm *et al*., 2016). These platforms were used to conduct the GWAS using as input the obtained data for the normalized relative primary root (PR) growth inhibition percentage in response to IAM treatment (PR length_Control_ – PR length_IAM_). A mixed linear model was performed to obtain candidate SNPs which were filtered both by minor allele frequency (MAF) ≥ 0.05 and a suggestive relaxed *p*-value criterion < 10^-5^ to consider possible IAM response-related genomic areas. Genes enclosed within a 10 kb-region around the identified SNPs were listed and ranked.

### Modelling of the ABA3 protein structures

The three-dimensional structures of the reference ABA3_Ref protein from Col-0 and the ABA3_Alt version containing all the common SNPs in the Arabidopsis accessions that showed reduced sensitivity towards IAM were modeled by using a homology-based approach. The 1.65 Å crystal structure of the cysteine desulfurase (SufS) from *Mycobacterium tuberculosis* [PDB: 8ODQ] (Elchennawi *et al*., 2023) and the 1.80 Å crystal structure of *Synechocystis* sp. PCC 6803 cysteine desulfurase (SufS) [PDB: 1T3I] (Tirupati *et al*., 2004) deposited in the Research Collaboratory for Structural Bioinformatics (RCSB) Protein Data Base were used as reference structures. The different structural models were generated by using both the Phyre^2^ protein fold recognition server (Kelley et al., 2015) and the I-TASSER protein structure prediction server (Yang et al., 2015). The structural comparison of the obtained models for ABA3_Ref and ABA3_Mod was performed using PyMOL v2.5.5 (https://pymol.org/) and the InterPro (https://www.ebi.ac.uk/interpro/) protein classification tool.

### Analysis of luciferase activity

For *in vivo* evaluation of changes in ABA signaling activity in response to treatment with IAM, bioluminescence measurements were performed using the pMAPKKK18::Luc^+^ reporter line. To this end, the reporter line was grown vertically for 10 days on 0.5× MS plates containing 1% (w/v) sucrose. The plants were then treated with a mock solution or a solution containing 20 µM IAM for 2 h, before monitoring luciferase activity using a cooled CCD camera (NightOwl II LB 983 NC-100; Berthold Technologies). To visualize the luciferase activity, plates were sprayed with 100 µM luciferin and imaged after an incubation time of 40 min.

### Confocal laser scanning microscopy

ABA signaling activities were also investigated in the roots of mock- and IAM-treated seedlings of ABA signaling reporter lines 6×ABRE_A::erGFP and 6×ABRE_R::erGFP using a Leica SP8 microscope with the Leica Application Suite (Las AF Lite) X software. As described above, the reporter plants were incubated for 2 h in a mock solution or in a solution containing either 20 µM IAM or 10 µM ABA. Thereafter, the green fluorescent protein (GFP) was excited at 488 nm using an Argon multiline laser and detected using a 494-596 nm broadband filter.

### RNA isolation and gene expression analysis by qRT-PCR

For the extraction of total RNA, 100 mg of plant tissue of ten days-old sterilely grown seedlings were harvested as previously described (Oñate-Sánchez & Vicente-Carbajosa, 2008). First strand synthesis was conducted using M-MLV reverse transcriptase and oligo(dT)_15_ primer, following the instructions of the manufacturer (Promega). Two nanograms of cDNA were used as template in each qRT-PCR. cDNA amplification was performed using the FastStart SYBR Green Master solution (Roche Diagnostics) and a Lightcycler 480 Real-Time PCR system (Roche Diagnostics), according to the supplier’s instructions. The relative transcript quantification was calculated employing the comparative 2^-ΔΔ*C*T^ method (Livak & Schmittgen, 2001). As reference genes, we used *APT1* (At1g27450) and *GAPC2* (At1g13440) (Czechowski *et al*., 2005; Jost *et al*., 2007). All experiments were carried out using biological triplicates. In addition, three technical replicates per biological replicate were analyzed. See **Supporting Information Table S4** for primer sequences.

### Comparative analysis of selected gene sets

The functional classification of DEGs was performed using the MapMan v3.6.0R1 software (Thimm *et al*., 2004), paying special attention to DEGs related with plant hormones. Furthermore, functional relationships between the DEGs were investigated employing the stringApp v2.0.3 (Doncheva *et al*., 2019), and EnrichmentMap v3.3.6 (Merico *et al*., 2010) in Cytoscape v3.10.1 (Shannon *et al*., 2003). To analyze the importance of the nodes in the inferred networks, the nodes with the highest degree of connectivity (k) and betweenness centrality (BC) were examined in closer detail.

### Statistical analysis

The statistical assessment of the data was performed using the JASP v0.18.3 software (https://jasp-stats.org/). Student’s *t*-test was employed to compare two means. Statistical differences of more than two means were determined by one-way ANOVA and Tukey’s post-hoc test for pairwise comparisons. Results were considered significant when the *p*-value < 0.05.

## Acknowledgements

The authors thank the NASC for providing Arabidopsis mutant seeds. Moreover, the authors are grateful to Jorge Lozano-Juste (Instituto de Biología Molecular y Celular de Plantas, Spain) for kindly sharing the pMAPKKK18-Luc^+^ reporter line with us. This research was funded by the Spanish Ministry of Science and Innovation (MICIN), grant BFU2017-82826-R and PID2020-119441RB-100 to S.P. and grant PID2022-136893NB-I00 to C.A.-B. from the MICIN/Agencia Estatal de Investigación of Spain (AEI)/10.13039/501100011033 and FEDER (EU). In addition, the authors acknowledge support by grant RED2022-134917-T funded by MICIN/AEI/10.13039/501100011033. J.M.-C. was supported by the ‘Severo Ochoa Program for Centers of Excellence in R&D’ from the AEI, grant SEV-2016-0672 (2017-2021) to the CBGP.

## Author Contributions

SP, JVC, JPA, CAB conceived and designed the research; JMC, POG, AGOV, IVL, APG, JVC and SP performed the research and analyzed the data; SP, JVC, JPA, and CAB were responsible for the acquisition of the required funding to perform the experiments and wrote and edited the manuscript. JMC and POG contributed equally. All authors have read and agreed to the published version of the manuscript.

## Short legends for Supporting Information

Supporting information Table S1: Metadata of the employed Arabidopsis thaliana accessions.

Supporting information Table S2: List of candidate genes obtained in the GWA study.

Supporting information Table S3: Transcriptional response to ABA and IAM treatments and Gene ontology (GO) enrichment analysis of the 206 DEGs responding to both ABA and IAM.

Supporting information Table S4: Primers used in this study.

Supporting information Figure S1: Secondary structure comparison of ABA3_Alt and ABA3_Ref.

Supporting information Figure S2: Transcriptional regulation of AMI1 and ABA3 by abiotic stress stimuli.

## References

Antoni R, Gonzalez-Guzman M, Rodriguez L, Peirats-Llobet M, Pizzio GA, Fernandez MA, De Winne N, De Jaeger G, Dietrich D, Bennett MJ, et al. 2013. PYRABACTIN RESISTANCE1-LIKE8 plays an important role for the regulation of abscisic acid signaling in root. Plant Physiology 161(2): 931–941.

Arteaga N, Savic M, Mendez-Vigo B, Fuster-Pons A, Torres-Perez R, Oliveros JC, Pico FX, Alonso-Blanco C. 2021. MYB transcription factors drive evolutionary innovations in Arabidopsis fruit trichome patterning. The Plant Cell 33(3): 548–565.

Barrett T, Wilhite SE, Ledoux P, Evangelista C, Kim IF, Tomashevsky M, Marshall KA, Phillippy KH, Sherman PM, Holko M, et al. 2012. NCBI GEO: archive for functional genomics data sets—update. Nucleic Acids Research 41(D1): D991–D995.

Bernhardt C, Lee MM, Gonzalez A, Zhang F, Lloyd A, Schiefelbein J. 2003. The bHLH genes *GLABRA3* (*GL3*) and *ENHANCER OF GLABRA3* (*EGL3*) specify epidermal cell fate in the *Arabidopsis* root. Development 130(26): 6431–6439.

Bhaskara GB, Nguyen TT, Verslues PE. 2012. Unique drought resistance functions of the *highly ABA-induced* clade A protein phosphatase 2Cs. Plant Physiology 160(1): 379– 395.

Bittner F, Oreb M, Mendel RR. 2001. ABA3 Is a Molybdenum Cofactor Sulfurase Required for Activation of Aldehyde Oxidase and Xanthine Dehydrogenase in *Arabidopsis thaliana*. Journal of Biological Chemistry 276(44): 40381–40384.

Bradford KJ, Trewavas AJ. 1994. Sensitivity Thresholds and Variable Time Scales in Plant Hormone Action. Plant Physiology 105(4): 1029–1036.

Brennan AC, Méndez-Vigo B, Haddioui A, Martínez-Zapater JM, Picó FX, Alonso-Blanco C. 2014. The genetic structure of Arabidopsis thaliana in the south-western Mediterranean range reveals a shared history between North Africa and southern Europe. BMC Plant Biology 14(1): 17.

Capriotti E, Fariselli P, Casadio R. 2005. I-Mutant2.0: predicting stability changes upon mutation from the protein sequence or structure. Nucleic Acids Research 33(Web Server issue): W306–310.

Chang TG, Zhu XG. 2017. Source-sink interaction: a century old concept under the light of modern molecular systems biology. Journal of Experimental Botany 68(16): 4417– 4431.

Czechowski T, Stitt M, Altmann T, Udvardi MK, Scheible WR. 2005. Genome-wide identification and testing of superior reference genes for transcript normalization in Arabidopsis. Plant Physiology 139(1): 5–17.

Davies PJ. 2010. Plant hormones. Biosynthesis, Signal Transduction, Action! Dordrecht, Boston, London: Springer Netherlands.

Depuydt S, Hardtke CS. 2011. Hormone signalling crosstalk in plant growth regulation. Current Biology 21(9): R365–373.

Doncheva NT, Morris JH, Gorodkin J, Jensen LJ. 2019. Cytoscape StringApp: Network Analysis and Visualization of Proteomics Data. Journal of Proteome Research 18(2): 623–632.

Elchennawi I, Carpentier P, Caux C, Ponge M, Ollagnier de Choudens S. 2023. Structural and Biochemical Characterization of *Mycobacterium tuberculosis* Zinc SufU-SufS Complex. Biomolecules 13(5): 732.

Escamez S, André D, Sztojka B, Bollhöner B, Hall H, Berthet B, Voß U, Lers A, Maizel A, Andersson M, et al. 2020. Cell Death in Cells Overlying Lateral Root Primordia Facilitates Organ Growth in *Arabidopsis*. Current Biology 30(3): 455–464.e457.

Finch-Savage WE, Leubner-Metzger G. 2006. Seed dormancy and the control of germination. New Phytologist 171(3): 501–523.

Fujii H, Verslues PE, Zhu JK. 2007. Identification of two protein kinases required for abscisic acid regulation of seed germination, root growth, and gene expression in *Arabidopsis*. The Plant Cell 19(2): 485–494.

García-Maquilón I, Rodriguez PL, Vaidya AS, Lozano-Juste J 2021. A LuciferaseLuciferase ReporterReportersAssay to Identify Chemical Activators of ABAAbscisic acid (ABA)Signaling. In: Hicks GR, Zhang C eds. Plant Chemical Genomics: Methods and Protocols. New York, NY: Springer US, 113–121.

Genomes Consortium. 2016. 1,135 Genomes Reveal the Global Pattern of Polymorphism in *Arabidopsis thaliana*. Cell 166(2): 481–491.

Ghassemian M, Nambara E, Cutler S, Kawaide H, Kamiya Y, McCourt P. 2000. Regulation of abscisic acid signaling by the ethylene response pathway in Arabidopsis. The Plant Cell 12(7): 1117–1126.

Goda H, Sasaki E, Akiyama K, Maruyama-Nakashita A, Nakabayashi K, Li W, Ogawa M, Yamauchi Y, Preston J, Aoki K, et al. 2008. The AtGenExpress hormone and chemical treatment data set: experimental design, data evaluation, model data analysis and data access. The Plant Journal 55(3): 526–542.

Gonzalez-Guzman M, Pizzio GA, Antoni R, Vera-Sirera F, Merilo E, Bassel GW, Fernández MA, Holdsworth MJ, Perez-Amador MA, Kollist H, et al. 2012. *Arabidopsis* PYR/PYL/RCAR receptors play a major role in quantitative regulation of stomatal aperture and transcriptional response to abscisic acid. The Plant Cell 24(6): 2483–2496.

Grimm DG, Roqueiro D, Salomé PA, Kleeberger S, Greshake B, Zhu W, Liu C, Lippert C, Stegle O, Schölkopf B, et al. 2016. easyGWAS: A Cloud-Based Platform for Comparing the Results of Genome-Wide Association Studies. The Plant Cell 29(1): 5–19.

Heidenreich T, Wollers S, Mendel RR, Bittner F. 2005. Characterization of the NifS-like Domain of ABA3 from *Arabidopsis thaliana* Provides Insight into the Mechanism of Molybdenum Cofactor Sulfuration. Journal of Biological Chemistry 280(6): 4213– 4218.

Holsteens K, De Jaegere I, Wynants A, Prinsen ELJ, Van de Poel B. 2022. Mild and severe salt stress responses are age-dependently regulated by abscisic acid in tomato. Frontiers in Plant Science 13: 982622.

Hsu P-K, Dubeaux G, Takahashi Y, Schroeder JI. 2021. Signaling mechanisms in abscisic acid-mediated stomatal closure. The Plant Journal 105(2): 307–321.

Jia W, Wang Y, Zhang S, Zhang J. 2002. Salt-stress-induced ABA accumulation is more sensitively triggered in roots than in shoots. Journal of Experimental Botany 53(378): 2201–2206.

Jost R, Berkowitz O, Masle J. 2007. Magnetic quantitative reverse transcription PCR: A high-throughput method for mRNA extraction and quantitative reverse transcription PCR. BioTechniques 43(2): 206–211.

Kasahara H. 2016. Current aspects of auxin biosynthesis in plants. Biosci Biotechnol Biochem 80(1): 34–42.

Knight H, Knight MR. 2001. Abiotic stress signalling pathways: specificity and cross-talk. Trends in Plant Science 6(6): 262–267.

Kuromori T, Seo M, Shinozaki K. 2018. ABA Transport and Plant Water Stress Responses. Trends in Plant Science 23(6): 513–522.

Léon-Kloosterziel KM, Gil MA, Ruijs GJ, Jacobsen SE, Olszewski NE, Schwartz SH, Zeevaart JAD, Koornneef M. 1996. Isolation and characterization of abscisic acid-deficient *Arabidopsis* mutants at two new loci. The Plant Journal 10(4): 655–661.

Liu X, Hou X. 2018. Antagonistic Regulation of ABA and GA in Metabolism and Signaling Pathways. Frontiers in Plant Science 9: 251.

Livak KJ, Schmittgen TD. 2001. Analysis of relative gene expression data using real-time quantitative PCR and the 2^-ΔΔ*C*^T Method. Methods 25(4): 402–408.

Maki H, Sakaoka S, Itaya T, Suzuki T, Mabuchi K, Amabe T, Suzuki N, Higashiyama T, Tada Y, Nakagawa T, et al. 2019. ANAC032 regulates root growth through the MYB30 gene regulatory network. Scientific Reports 9(1): 11358.

Marcer A, Vidigal DS, James PMA, Fortin MJ, Méndez-Vigo B, Hilhorst HWM, Bentsink L, Alonso-Blanco C, Picó FX. 2018. Temperature fine-tunes Mediterranean *Arabidopsis thaliana* life-cycle phenology geographically. Plant Biology (Stuttg) 20 Suppl 1: 148–156.

Merico D, Isserlin R, Stueker O, Emili A, Bader GD. 2010. Enrichment Map: A Network-Based Method for Gene Set Enrichment Visualization and Interpretation. PLoS One 5(11): e13984.

Motte H, Vanneste S, Beeckman T. 2019. Molecular and Environmental Regulation of Root Development. Annual Review of Plant Biology 70: 465–488.

Moya-Cuevas J, Pérez-Alonso MM, Ortiz-García P, Pollmann S. 2021. Beyond the Usual Suspects: Physiological Roles of the Arabidopsis Amidase Signature (AS) Superfamily Members in Plant Growth Processes and Stress Responses. Biomolecules 11(8): 1207.

Murashige T, Skoog F. 1962. A revised medium for rapid growth and bio assays with tobacco tissue cultures. Physiologia Plantarum 15(3): 473–497.

Nambara E, Marion-Poll A. 2003. ABA action and interactions in seeds. Trends in Plant Science 8(5): 213–217.

Nambara E, Suzuki M, Abrams S, McCarty DR, Kamiya Y, McCourt P. 2002. A Screen for Genes That Function in Abscisic Acid Signaling in *Arabidopsis thaliana*. Genetics 161(3): 1247–1255.

Nemoto K, Hara M, Suzuki M, Seki H, Muranaka T, Mano Y. 2009. The *NtAMI1* gene functions in cell division of tobacco BY-2 cells in the presence of indole-3-acetamide. FEBS Letters 583(2): 487–492.

Oñate-Sánchez L, Vicente-Carbajosa J. 2008. DNA-free RNA isolation protocols for *Arabidopsis thaliana*, including seeds and siliques. BMC Research Notes 1: 93.

Ortiz-García P, González Ortega-Villaizán A, Onejeme FC, Müller M, Pollmann S. 2023. Do Opposites Attract? Auxin-Abscisic Acid Crosstalk: New Perspectives. International Journal of Molecular Sciences 24(4).

Ortiz-García P, Pérez-Alonso MM, González Ortega-Villaizan A, Sánchez-Parra B, Ludwig-Müller J, Wilkinson MD, Pollmann S. 2022. The Indole-3-Acetamide-Induced Arabidopsis Transcription Factor MYB74 Decreases Plant Growth and Contributes to the Control of Osmotic Stress Responses. Frontiers in Plant Science 13: 928386.

Park S-Y, Fung P, Nishimura N, Jensen DR, Fujii H, Zhao Y, Lumba S, Santiago J, Rodrigues A, Chow T-fF, et al. 2009. Abscisic Acid Inhibits Type 2C Protein Phosphatases via the PYR/PYL Family of START Proteins. Science 324(5930): 1068– 1071.

Pérez-Alonso MM, Ortiz-García P, Moya-Cuevas J, Lehmann T, Sánchez-Parra B, Björk RG, Karim S, Amirjani MR, Aronsson H, Wilkinson MD, et al. 2021. Endogenous indole-3-acetamide levels contribute to the crosstalk between auxin and abscisic acid, and trigger plant stress responses in *Arabidopsis thaliana*. Journal of Experimental Botany 72(2): 459–475.

Picó FX, Méndez-Vigo B, Martínez-Zapater JM, Alonso-Blanco C. 2008. Natural genetic variation of *Arabidopsis thaliana* is geographically structured in the Iberian peninsula. Genetics 180(2): 1009–1021.

Pires DE, Ascher DB, Blundell TL. 2014. DUET: a server for predicting effects of mutations on protein stability using an integrated computational approach. Nucleic Acids Research 42(Web Server issue): W314–319.

Pollmann S, Müller A, Weiler EW. 2006. Many roads lead to “auxin”: of nitrilases, synthases, and amidases. Plant Biology (Stuttg) 8(3): 326–333.

Pollmann S, Neu D, Weiler EW. 2003. Molecular cloning and characterization of an amidase from *Arabidopsis thaliana* capable of converting indole-3-acetamide into the plant growth hormone, indole-3-acetic acid. Phytochemistry 62(3): 293–300.

Ré DA, Capella M, Bonaventure G, Chan RL. 2014. Arabidopsis AtHB7 and AtHB12evolved divergently to fine tune processes associated with growth and responses to water stress. BMC Plant Biology 14(1): 150.

Rieu I, Eriksson S, Powers SJ, Gong F, Griffiths J, Woolley L, Benlloch R, Nilsson O, Thomas SG, Hedden P, et al. 2008. Genetic Analysis Reveals That C_19_-GA 2-Oxidation Is a Major Gibberellin Inactivation Pathway in *Arabidopsis* The Plant Cell 20(9): 2420–2436.

Rizza A, Tang B, Stanley CE, Grossmann G, Owen MR, Band LR, Jones AM. 2021. Differential biosynthesis and cellular permeability explain longitudinal gibberellin gradients in growing roots. Proceedings of the National Academy of Sciences USA 118(8): 154498.

Robe K, Barberon M. 2023. Nutrient carriers at the heart of plant nutrition and sensing. Current Opinion in Plant Biology 74: 102376.

Rubio S, Rodrigues A, Saez A, Dizon MB, Galle A, Kim T-H, Santiago J, Flexas J, Schroeder JI, Rodriguez PL. 2009. Triple Loss of Function of Protein Phosphatases Type 2C Leads to Partial Constitutive Response to Endogenous Abscisic Acid. Plant Physiology 150(3): 1345–1355.

Sánchez-Parra B, Frerigmann H, Pérez-Alonso MM, Carrasco-Loba V, Jost R, Hentrich M, Pollmann S. 2014. Characterization of Four Bifunctional Plant IAM/PAM-Amidohydrolases Capable of Contributing to Auxin Biosynthesis. Plants (Basel) 3(3): 324–347.

Sánchez-Parra B, Pérez-Alonso MM, Ortiz-García P, Moya-Cuevas J, Hentrich M, Pollmann S. 2021. Accumulation of the Auxin Precursor Indole-3-Acetamide Curtails Growth through the Repression of Ribosome-Biogenesis and Development-Related Transcriptional Networks. International Journal of Molecular Sciences 22(4): 2040.

Santner A, Calderon-Villalobos LIA, Estelle M. 2009. Plant hormones are versatile chemical regulators of plant growth. Nature Chemical Biology 5(5): 301–307.

Schindelin J, Arganda-Carreras I, Frise E, Kaynig V, Longair M, Pietzsch T, Preibisch S, Rueden C, Saalfeld S, Schmid B, et al. 2012. Fiji: an open-source platform for biological-image analysis. Nature Methods 9(7): 676–682.

Schwarz G, Mendel RR, Ribbe MW. 2009. Molybdenum cofactors, enzymes and pathways. Nature 460(7257): 839–847.

Seo M, Peeters AJM, Koiwai H, Oritani T, Marion-Poll A, Zeevaart JAD, Koornneef M, Kamiya Y, Koshiba T. 2000. The *Arabidopsis* aldehyde oxidase 3 (*AAO3*) gene product catalyzes the final step in abscisic acid biosynthesis in leaves. Proceedigns of the National Academy of Sciences USA 97(23): 12908–12913.

Seren Ü, Vilhjálmsson BJ, Horton MW, Meng D, Forai P, Huang YS, Long Q, Segura V, Nordborg M. 2012. GWAPP: A Web Application for Genome-Wide Association Mapping in Arabidopsis The Plant Cell 24(12): 4793–4805.

Shannon P, Markiel A, Ozier O, Baliga NS, Wang JT, Ramage D, Amin N, Schwikowski B, Ideker T. 2003. Cytoscape: A Software Environment for Integrated Models of Biomolecular Interaction Networks. Genome Research 13(11): 2498–2504.

Shinkawa R, Morishita A, Amikura K, Machida R, Murakawa H, Kuchitsu K, Ishikawa M. 2013. Abscisic acid induced freezing tolerance in chilling-sensitive suspension cultures and seedlings of rice. BMC Research Notes 6(1): 351.

Sugawara S, Hishiyama S, Jikumaru Y, Hanada A, Nishimura T, Koshiba T, Zhao Y, Kamiya Y, Kasahara H. 2009. Biochemical analyses of indole-3-acetaldoxime-dependent auxin biosynthesis in *Arabidopsis*. Proceedigns of the National Academy of Sciences USA 106(13): 5430–5435.

Sukiran NL, Ma JC, Ma H, Su Z. 2019. ANAC019 is required for recovery of reproductive development under drought stress in *Arabidopsis*. Plant Molecular Biology 99(1): 161–174.

Tabas-Madrid D, Méndez-Vigo B, Arteaga N, Marcer A, Pascual-Montano A, Weigel D, Xavier Picó F, Alonso-Blanco C. 2018. Genome-wide signatures of flowering adaptation to climate temperature: Regional analyses in a highly diverse native range of *Arabidopsis thaliana*. Plant, Cell & Environment 41(8): 1806–1820.

Takatsuka H, Umeda M. 2014. Hormonal control of cell division and elongation along differentiation trajectories in roots. Journal of Experimental Botany 65(10): 2633– 2643.

Teale WD, Paponov IA, Palme K. 2006. Auxin in action: signalling, transport and the control of plant growth and development. Nature Reviews Molecular Cell Biology 7(11): 847–859.

Thimm O, Bläsing O, Gibon Y, Nagel A, Meyer S, Krüger P, Selbig J, Müller LA, Rhee SY, Stitt M. 2004. MAPMAN: a user-driven tool to display genomics data sets onto diagrams of metabolic pathways and other biological processes. The Plant Journal 37(6): 914–939.

Thole JM, Beisner ER, Liu J, Venkova SV, Strader LC. 2014. Abscisic acid regulates root elongation through the activities of auxin and ethylene in *Arabidopsis thaliana*. G3 (Bethesda) 4(7): 1259–1274.

Tirupati B, Vey JL, Drennan CL, Bollinger JM. 2004. Kinetic and Structural Characterization of Slr0077/SufS, the Essential Cysteine Desulfurase from *Synechocystis* sp. PCC 6803. Biochemistry 43(38): 12210–12219.

Vidigal DS, Marques AC, Willems LA, Buijs G, Méndez-Vigo B, Hilhorst HW, Bentsink L, Picó FX, Alonso-Blanco C. 2016. Altitudinal and climatic associations of seed dormancy and flowering traits evidence adaptation of annual life cycle timing in *Arabidopsis thaliana*. *Plant*, Cell & Environment 39(8): 1737–1748.

Waese J, Fan J, Pasha A, Yu H, Fucile G, Shi R, Cumming M, Kelley LA, Sternberg MJ, Krishnakumar V, et al. 2017. ePlant: Visualizing and Exploring Multiple Levels of Data for Hypothesis Generation in Plant Biology. Plant Cell 29(8): 1806–1821.

Wang S, Cao L, Willick IR, Wang H, Tanino KK. 2022. Arabidopsis Ubiquitin-Conjugating Enzymes UBC4, UBC5, and UBC6 Have Major Functions in Sugar Metabolism and Leaf Senescence. International Journal of Molecular Sciences 23(19): 11143.

Wolters H, Jürgens G. 2009. Survival of the flexible: hormonal growth control and adaptation in plant development. Nature Reviews Genetics 10(5): 305–317.

Wong C, Alabadí D, Blázquez MA. 2023. Spatial regulation of plant hormone action. Journal of Experimental Botany 74(19): 6089–6103.

Woodward AW, Bartel B. 2005. Auxin: regulation, action, and interaction. Annals of Botany 95(5): 707–735.

Wu R, Duan L, Pruneda-Paz JL, Oh DH, Pound M, Kay S, Dinneny JR. 2018. The 6xABRE Synthetic Promoter Enables the Spatiotemporal Analysis of ABA-Mediated Transcriptional Regulation. Plant Physiology 177(4): 1650–1665.

Xu Y, Prunet N, Gan ES, Wang Y, Stewart D, Wellmer F, Huang J, Yamaguchi N, Tatsumi Y, Kojima M, et al. 2018. SUPERMAN regulates floral whorl boundaries through control of auxin biosynthesis. The EMBO Journal 37(11): e97499.

Yang J, Zhang Y. 2015. I-TASSER server: new development for protein structure and function predictions. Nucleic Acids Research 43(W1): W174–181.

Yang L, Zhang J, He J, Qin Y, Hua D, Duan Y, Chen Z, Gong Z. 2014. ABA-mediated ROS in mitochondria regulate root meristem activity by controlling *PLETHORA* expression in *Arabidopsis*. PLoS Genetics 10(12): e1004791.

Yoshida T, Christmann A, Yamaguchi-Shinozaki K, Grill E, Fernie AR. 2019. Revisiting the Basal Role of ABA - Roles Outside of Stress. Trends in Plant Science 24(7): 625– 635.

Yoshida T, Fernie AR. 2023. Hormonal regulation of plant primary metabolism under drought. Journal of Experimental Botany 75(6): 1714–1725.

Zhao Y. 2010. Auxin biosynthesis and its role in plant development. Annual Review of Plant Biology 61: 49–64.

Zheng Z, Xu X, Crosley RA, Greenwalt SA, Sun Y, Blakeslee B, Wang L, Ni W, Sopko MS, Yao C, et al. 2010. The protein kinase SnRK2.6 mediates the regulation of sucrose metabolism and plant growth in Arabidopsis. Plant Physiology 153(1): 99–113.

