## Supplementary figures and images for "Identification of a novel link connecting indole-3-acetamide with abscisic acid biosynthesis and signaling"

### Supporting Information Figure 1

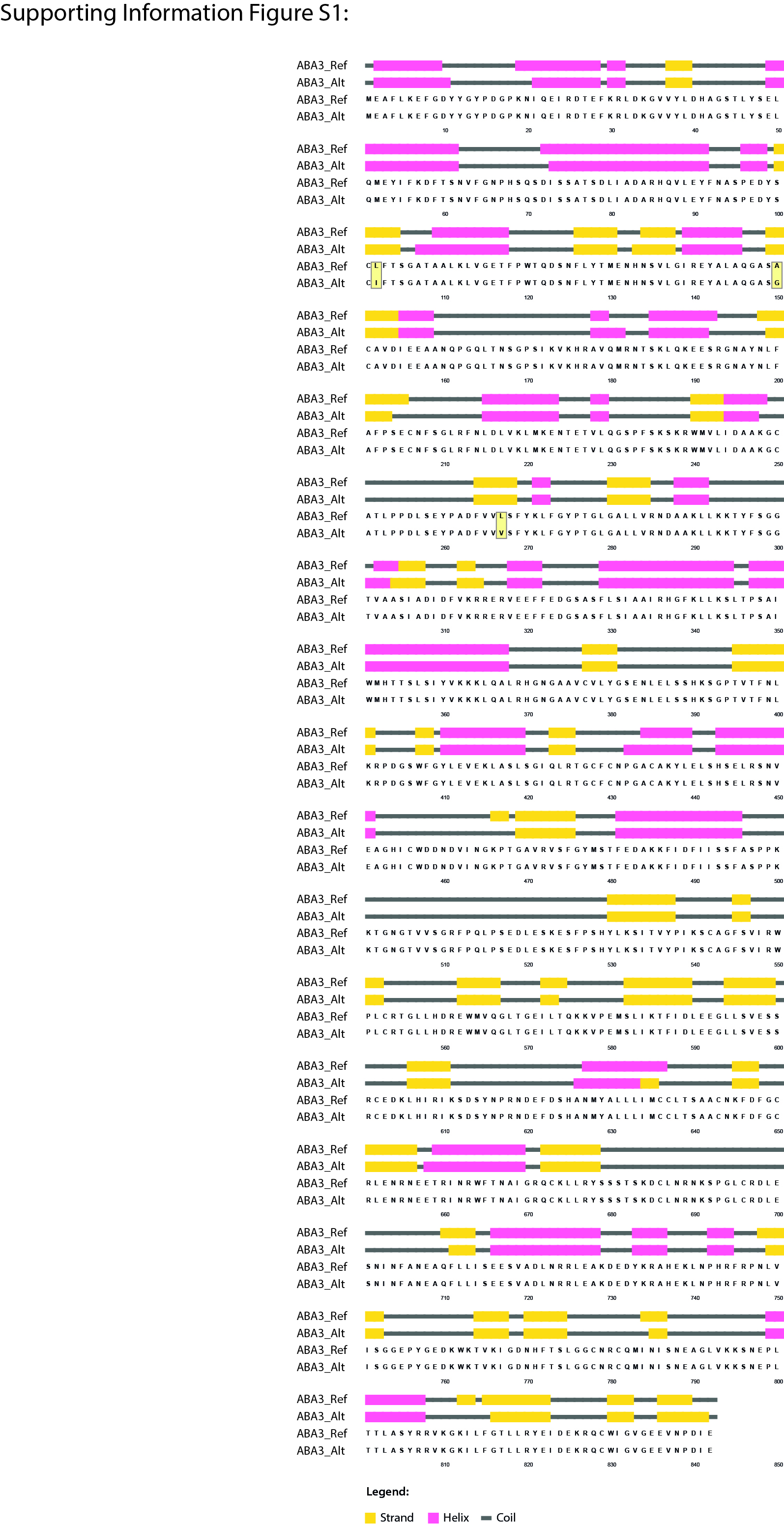

### Supporting Information Figure 2

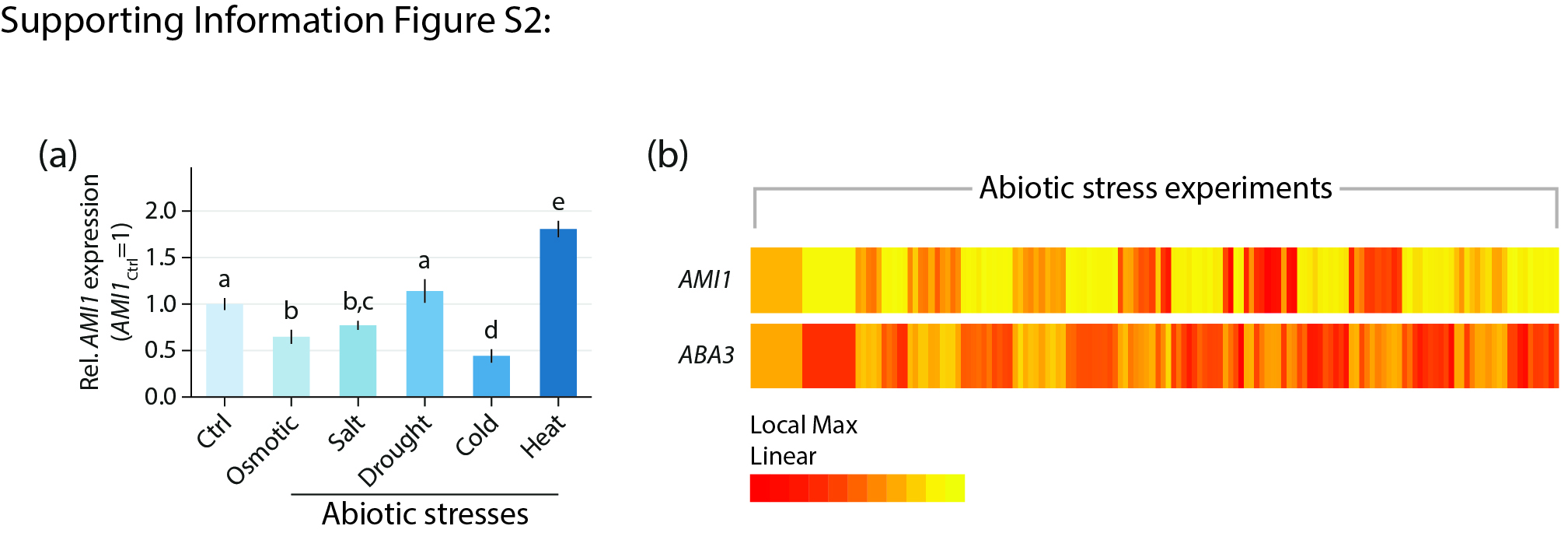
